# Electrophysiological dynamics of empathy for self- and other-related pain

**DOI:** 10.64898/2026.11.12.765918

**Authors:** Parisa Rahnama Lashkami, Soroosh Golbabaei, Shahriar Gharibzadeh, Khatereh Borhani

**Affiliations:** Institute for Cognitive and Brain Sciences, Shahid Beheshti University, Tehran, Iran; Department of Psychiatry and Psychotherapy, Jena University Hospital, Jena, Germany; German Center for Mental Health (DZPG), partner site Halle-Jena-Magdeburg, Germany

**Keywords:** empathy, painful facial expressions, self–other distinction, shared representations, Time-frequency Analysis, EEG, ERP

## Abstract

Empathy requires the ability to generate shared representations of the self and others while maintaining a distinction between them. However, it remains unclear whether, and to what extent, the neural processes involved in sharing and understanding others’ pain overlap with those involved in processing one’s own pain, particularly with respect to their temporal dynamics. Therefore, the present study aimed to investigate the neural mechanisms underlying empathic processing of self- and other-related painful facial expressions. To this end, EEG signals were recorded from 42 female participants while they viewed self- and other-related painful and neutral expressions. During the early stage of processing, the difference in N2 amplitude between self-related and other-related stimuli was smaller in the painful than in the neutral condition. Nevertheless, self-related painful stimuli continued to elicit more negative N2 amplitudes and greater Alpha/Mu suppression than other-related painful stimuli. In contrast, during the late stage of empathic processing, no distinction was observed between self- and other-related painful stimuli. Instead, painful expressions, regardless of whether they were self- or other-related, elicited larger late positive potential (LPP) amplitudes. These findings suggest that empathic processing is characterized by an early self–other distinction followed by a later convergence in the neural representation of self- and other-related pain.

## Introduction

Empathy is a multidimensional psychological construct that enables individuals to understand and share the emotional and affective states of others while maintaining a distinction between the experiences of the self and other (Decety & Lamm, 2006). It is commonly conceptualized as comprising affective and cognitive components. Affective empathy refers to the tendency to experience and share the emotions of others, whereas cognitive empathy involves understanding and interpreting the emotions, thoughts, and perspectives of others (Davis, 1983, 2006; Decety & Jackson, 2004; Dvash & Shamay-Tsoory, 2014). Although these components are distinguishable, empathic responding depends on their coordinated operation and, critically, on the ability to relate another person’s experience to one’s own without confusing the two.

One mechanism proposed to support this process is shared representation. Observing the states or actions of others can engage neural responses similar to those evoked during the first-hand experience of the same states or actions (Decety & Hodges, 2006; Knoblich & Flach, 2003; Lamm et al., 2011). Thus, understanding another person’s experience may be grounded, at least in part, in representations derived from one’s own experiences. However, representational overlap alone is insufficient for empathy. Empathic processing also requires maintaining self-other distinction the capacity to differentiate between self-related states and those attributed to another person (Decety & Sommerville, 2003; Jackson et al., 2006; Jackson et al., 2005). The balance between shared representations and self-other distinction is therefore considered a fundamental prerequisite for empathy (Davis, 1983; Lamm et al., 2016; Zaki et al., 2007). This balance may be particularly relevant to empathy for pain. Comparing responses to empathy evoking facial expressions associated with oneself and with another person provide an opportunity to examine the extent to which empathic processing relies on shared representations while remaining sensitive to the distinction between self and other. Importantly, self–other distinction is unlikely to constitute a single, static operation and is instead suggested to be a dynamic, time-dependent process that is likely to emerge progressively across different stages of empathic processing (Lamm et al., 2016; Tajadura-Jiménez et al., 2012). Thus, determining when neural responses to one’s own and another person’s pain converge and diverge may provide insight into the temporal development of shared and self-specific processes during empathy.

Electroencephalography (EEG), and event-related potentials (ERPs) are particularly well suited for addressing this question because of its high temporal resolution. Previous ERP studies of empathy for pain, many of which have used images depicting painful stimulation to body parts, have implicated both early components such as N1 and N2, and late components such as the late positive potential, with significant effects most consistently reported for the N2 and LPP (Liu et al., 2023; Liu et al., 2019; Suzuki et al., 2015). Some studies have linked the early N1 and N2 components to rapid, automatic aspects of empathy, including processing of others’ emotional states, affective responsiveness, and the initial allocation of attention to cognitively or emotionally salient stimuli (Carretié et al., 2004; Cui et al., 2016; Fan & Han, 2008; Liu et al., 2019; Meng et al., 2019; Zhu et al., 2019). In contrast, the LPP has been linked to more sustained and elaborative processing of emotionally and motivationally relevant stimuli (Carretié et al., 2004; Fan & Han, 2008; Hajcak & Foti, 2020; Li et al., 2019; Olofsson et al., 2008; Sun et al., 2017). Studies have reported larger LPP amplitudes in response to painful than neutral stimuli (Branchadell et al., 2024; Liao et al., 2021; Yang et al., 2024). Together, these findings suggest that ERP components can provide complementary information about how pain-related information is processed across different temporal stages.

However, phase-locked ERP components alone can provide only a partial characterization of the electrophysiological dynamics associated with empathy, as they primarily reflect stimulus-synchronized activity and do not encompass induced oscillations that are not phase-locked to the stimulus (Gomarus et al., 2006; Lechinger et al., 2016). Time-frequency analyses can complement ERPs by quantifying stimulus-related changes in oscillatory activity, including event-related desynchronization/synchronization (ERD/ERS) that need not be phase-locked to the stimulus (Fabi & Leuthold, 2017). Of particular relevance is activity in the Alpha/Mu frequency range, within the 8–13 Hz, especially over sensorimotor regions, particularly at the central electrodes C3, Cz, and C4 (Grice-Jackson et al., 2017; Woodruff et al., 2011). Suppression of Alpha/mu rhythm has been associated with both first-hand and observed sensory experiences, as well as processing of social and pain-related information (Coll et al., 2015; Fabi & Leuthold, 2017; Ferguson et al., 2024; Gaetz & Cheyne, 2006; Oberman & Ramachandran, 2007; Perry et al., 2010; Riečanský et al., 2020). These findings have made Alpha/mu suppression a potentially informative index of the degree to which observed experiences engage representations related to one’s own sensorimotor states. Importantly, Alpha/mu activity may also be sensitive to self-other attribution. Riečanský et al. (2020), for example, manipulated the degree to which an observed hand experiencing pain was attributed to the participant and found greater the Alpha/mu-rhythm suppression with increasing self-attribution. This finding suggests that the electrophysiological response to observed pain is influenced not only by whether an event is painful, but also by the extent to which that experience is represented as related to the self. Whether comparable self-other differences emerge when pain is conveyed through facial expressions, however remains unclear. Moreover, because oscillatory response evolves over time, remaining Alpha/mu activity across different temporal windows may help clarify when shared versus self-related representations become apparent during pain processing and empathy.

Despite extensive research on the electrophysiological correlates of empathy for pain, previous studies have mainly examined responses to pain experienced by others. Relatively little is known about how neural responses to another person’s painful facial expression compare directly with responses to one’s own painful facial expression within the same paradigm. Such a comparison is particularly informative for distinguishing neural processes related to the detection and evaluation of pain from those related to self-other representation.

Accordingly, the present study investigated the temporal electrophysiological dynamics associated with viewing painful and neutral facial expressions of the self and an unfamiliar other. Participants viewed painful and neutral facial expressions belonging either to themselves or to another individual. We examined early ERP components (N1 and N2), the later LPP, and alpha/mu-band power across early and late time windows. By combining ERP and time– frequency measures, the study sought to characterize when neural responses to pain are shared across self and other and when they become differentiated as a function of self–other identity.

Because social understanding is partly grounded in self-related representations, individuals may rely on their own experiences and affective states as an initial reference when interpreting others’ emotions and mental states (Lamm et al., 2011). Based on this, it was expected that, during the early stages of empathic processing, a distinction would emerge between the processing of self-and other-related painful stimuli. In other words, individuals initially show stronger electrophysiological responses to self-related painful stimuli as a means of understanding others’ pain. Conversely, during the later stages of empathic processing, the painfulness of the stimulus, irrespective of the self–other distinction, was expected to be the primary determinant of the electrophysiological empathic response.

To achieve this objective, electroencephalography (EEG) data were recorded from female participants while they viewed self-and other-related painful and neutral facial expressions. Because previous studies have shown that gender may influence empathic processing (McDonald & Kanske, 2023; Mestre et al., 2009), this study included only female participants to control for gender. Accordingly, the present study hypothesized that: (i) Self-related painful stimuli would elicit a more negative N2 amplitude than other-related painful stimuli; and (ii) LPP amplitude would not differ significantly between self-related and other-related painful stimuli; and (iii) During the early stage of processing, Alpha/Mu rhythm suppression would be greater for self-related painful stimuli than for other-related painful stimuli.

## Methods

### Participants

The required sample size was determined using G*Power 3.1 software (Faul et al., 2007). Assuming a within-subjects design, an alpha level of 0.05, a power of 0.8, and medium effect size (*f* = 0.25) based on previous empathy studies (Chen et al., 2023; Wang et al., 2023) the minimum required sample was 24 participants. To account for potential exclusions and dropouts we recruited 43 female participants. One participant was excluded because of a self-reported diagnosis of a psychological disorder, resulting in a final sample of 42 participants (*M*_age_= 20.21, *SD*= 3.32, range= 18-34). All participants were native Persian speakers, had normal or corrected-to-normal vision, and no history of psychological disorders. The study was approved by the Ethical Research Committee of Shahid Beheshti University and was conducted in accordance with ethical principles of the Declaration of Helsinki. Written informed consent was obtained from all participants prior to participation.

### Stimuli

The stimuli used in this experiment consisted of photographs of each participant’s own face and the face of an unfamiliar female individual, each displaying painful and neutral expressions. The same unfamiliar female individual was used as the other-related stimulus for all participants. She was 27 years old at the time of photography and was not enrolled as a study participant; rather, she was photographed solely to produce the other-related images used as experimental stimuli. Her photographs were taken during the same period as those of the other participants. For each participant, five photographs were taken in the painful condition and five in the neutral condition, with images captured from five different head orientations (upward, downward, left, right, and straight ahead) to introduce visual variability and reduce stimulus repetition. The same procedure was applied to the unfamiliar female individual. To elicit pain-related facial expressions, participants received instructions regarding the characteristic facial movements associated with painful expression. In accordance with the Facial Action Coding System (FACS; Ekman & Friesen,1978), pain-related facial expressions can involve brow lowering, cheek raising, lid tightening, lip stretching, nose wrinkling, nasolabial deepening, lip corner pulling, lips parting, upper lip raising, jaw dropping, lid drooping, eye closure, and blinking (Craig et al., 1992). Participants were subsequently shown examples of painful facial expressions and instructed to imagine themselves enduring pain and to reproduce the corresponding facial expression. For each participant, the final stimulus set comprised 20 images: 10 self-related images (five painful and five neutral) and 10 other-related images (five painful and five neutral). All images were standardized to a size of 530 × 600 pixels and presented centrally against a black background.

### Passive Viewing of Self and Other Painful Facial Expressions Task

During EEG recording, participants completed the Passive Viewing of Self and Other Painful Expressions Task, which was designed using PsychoPy (2024.2.4; Peirce,2007). The paradigm used in this task was adapted from previous ERP studies investigating empathy for pain (Chen et al., 2023; Liu et al., 2023; Pan et al., 2023). During the task, participants completed a total of 800 trials across four blocks. Each block consisted of 200 trials. Within each block, there were 50 trials related to the participant’s painful face, 50 trials related to another person’s painful face, 50 trials related to the participant’s neutral face, and 50 trials related to another person’s neutral face. Each picture in each block was repeated ten times. During the recording, each trial began with a fixation cross presented for 500 ms, followed by a picture displayed for 1000 ms. Afterward, a blank screen appeared for 500 ms, followed by a question (“How intense is the pain expressed by the person in the picture?”) which was displayed for 3000 ms. Participants had to respond on a 7-point scale (1 = not at all, 7 = extremely). At the end of each block, a question (“Was the previous picture of you or someone else?”) was presented to check the participant’s attention. Participants had 5 s to respond by pressing key 1 (me) or key 2 (someone else). The inter-trial interval was 1000 ms. There was a 30 s rest period between each block. After the rest period, participants pressed the space key to begin the next block. Before the task began, participants completed two practice trials to become familiar with the procedure.

### Empathy Task

In line with previous studies (Golbabaei & Borhani, 2024; Golbabaei, Hosseinnezhad, et al., 2026; Golbabaei, Walter, et al., 2026; Pfabigan et al., 2015) the empathy task for assessing cognitive and affective components of empathy was created in Psychopy (2024.2.4; Peirce,2007) and used. Each trial began with presentation of a fixation cross for 500 ms. Subsequently, participants viewed a picture depicting either self or another person, shown in either painful or neutral condition, alongside a series of three questions. These pictures were the same as the ones used in the “passive viewing of self and other painful expression task”. First, they rated the level of pain perceived in the picture (1 = not at all, 7 = extremely). Second, they evaluated the extent of unpleasant emotion expressed by the face in the picture (1 = not at all, 7 = extremely); which indexed cognitive empathy. Finally, they rated the degree of unpleasant feeling they personally experienced while observing the picture (1 = not at all, 7 = extremely); which indexed affective empathy. After answering each question, a blank screen was presented for 500 ms, and the next question was then presented simultaneously with the picture.

### Procedure

Participants attended the laboratory for two separate sessions conducted on different days. During the first session, ten photographs of each participant were taken to create the experimental stimuli. In the second session, participants returned to the laboratory for EEG recording. They were seated comfortably in a quiet room approximately 80 cm away from the monitor screen. An EEG cap was fitted, and conductive gel was applied to ensure adequate signal quality. Following EEG preparation, participants completed the Passive Viewing of Self and Other Painful Facial Expression Task while EEG activity was continuously recorded. They were instructed to follow the task instructions carefully and to avoid unnecessary movements during the signal acquisition. This task lasted approximately 60 minutes. After completion, the EEG cap and electrodes were removed. Following a short break, participants proceeded to an Empathy Task without any time constraints. This Task lasted about 10 minutes.

### EEG Recording and Preprocessing

EEG was recorded using the Liv system (Liv Intelligent Technology, Iran) with 64 Ag/AgCl electrodes, positioned according to the international 10-20 system. Conductive gel was applied to reduce electrode-scalp impedance, which was kept below 5kΩ throughout the recording. The earlobes were used as an online reference. EEG signals were recorded at a sampling frequency of 250 Hz. EEG data were preprocessed using the EEGLAB toolbox (Delorme & Makeig, 2004) in MATLAB R2021b. The data were band-pass filtered between 0.5 and 30 Hz band-pass filter and subsequently re-referenced to the average of all channels.

Independent Component Analysis (ICA) was performed to identify and remove components associated with physiological and non-neural artifacts, including eye movements, muscle contractions, heartbeat, and channel noise (Jung et al., 2001). In addition, the EEG data were visually inspected to identify and remove residual artifacts and noisy segments.

#### ERP analysis

For each trial, EEG data were segmented into epochs extending from 200 ms before to 1000 ms after stimuli onset. Each epoch was baseline-corrected using a 200 ms pre-stimulus interval. Trials in which the amplitude exceeded ±100 μV at any time point within the epoch in any channel were considered artifact-contaminated and removed from further analysis. In total, approximately 1% of trials were removed in each condition (Other-Neutral, Other-Painful, Self-Neutral, Self-Painful). Based on the inspection of scalp topography and previous literature (Chen et al., 2024; Li et al., 2020; Li et al., 2022; Liao et al., 2021; Miedzobrodzka et al., 2023), specific electrodes and time windows were selected for each ERP component. The N1 component was quantified at F1, Fz, F2, FC1, FCz, and FC2 in the time window of 130-190 ms post-stimulus presentation. The N2 component was quantified at P7, P8, PO7, and PO8 in the time window of 170-290 ms, and the LPP was quantified at CP1, CPz, CP2, P1, Pz, P2, PO3, POz, and PO4 in the time window of 340-800 ms.

#### Time-frequency Analysis of Alpha/Mu Oscillations

EEG data were initially epoched from −1000 to +1000 ms relative to the onset of picture presentation. Time–frequency decomposition was performed using the Short-Time Fourier Transform (STFT), with a fixed 64-sample Hanning window (equivalent to 256 ms), and 50% overlap between consecutive windows. Spectral power was estimated across the 1–30 Hz frequency range. The analysis was conducted in the Alpha/Mu band. Consistent with previous studies (Chen et al., 2012; Cheng et al., 2014; Grice-Jackson et al., 2017; Woodruff et al., 2011), Alpha/Mu activity was quantified by averaging the spectral activity across the sensorimotor electrodes C3, Cz, and C4. The baseline spectral power at each frequency was defined as the average power during the prestimulus interval from −500 to 0 ms. Baseline correction was performed by normalizing the spectral power at each time–frequency point to the mean baseline power at the corresponding frequency. The resulting values were then converted to decibels (dB). The Event-Related Spectral Perturbation (ERSP) values were calculated using the following equation:

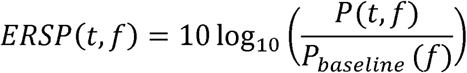

Alpha/Mu ERSP was calculated by averaging the ERSP values within the Alpha/Mu frequency band from 8 to 13 Hz separately for the early time window from 0 to 500 ms and the late time window from 500 to 1000 ms.

### Statistical Analysis

Statistical analyses were performed using R with the packages rstatix, tidyverse, and ggpubr. To examine the effects of Stimulus type (Self vs. Other) and Facial expression (Painful vs. Neutral) on ERP component amplitudes a 2 × 2 repeated-measures analysis of variance (ANOVA) was conducted. For, Alpha/Mu rhythm oscillatory changes, a 2 × 2 × 2 repeated-measures ANOVAs was performed with Stimulus type, Facial expression, and Time window (Early vs. Late) as within-subject factors. When a significant interaction effect was observed, post hoc comparisons were performed, and Bonferroni correction was applied to control for multiple comparisons.

## Results

### Manipulation Check

Paired-samples t-tests showed that pain ratings were significantly higher for painful than for neutral images in both the self condition, *t* (41) = 25.203, *p* < 0.001 (*M* _neutral_ = 1.143 ± 0.281; *M* _painful_ = 4.417 ± 0.925), and the other condition, *t* (41) = 28.139, *p* < 0.001 (*M* _neutral_ = 1.143 ± 0.250; *M* _painful_ = 4.541 ± 0.883).

### Behavioral results

A significant interaction of Stimulus type and Facial expression was observed for affective empathy ratings, *F* (1,41) = 7.314, *p* = 0.010, η_p_^2^ = 0.151. Post-hoc comparisons showed that, in the Painful condition, affective empathy ratings were significantly higher for other (4.705 ± 1.055) than for the self (4.376 ± 1.114), *t* (41) = 2.538, *p_corr_* = 0.030, *p* _=_ 0.015. In contrast, no significant difference between Self (1.371 ± 0.544) and Other (1.271 ± 0.533) was observed in the Neutral condition, *t* (41) = −1.527, *p_corr_* = 0.269, *p* _=_ 0.134. The main effect of Facial expression was significant, *F* (1,41) = 660.766; *p* < 0.001, η_p_^2^ = 0.942, with higher affective empathy ratings for Painful stimuli (4.540 ± 1.091) than for Neutral stimuli (1.321 ± 0.538). However, the main effect of Stimulus type was not significant, *F* (1,41) = 3.076, *p* = 0.087, η_p_^2^ = 0.070.

Similarly, a significant interaction of Stimulus type and Facial expression was observed for cognitive empathy ratings, *F* (1,41) = 13.642, *p* < 0.001, η_p_^2^ = 0.25. Post-hoc comparisons showed that, in the Painful condition, cognitive empathy ratings were significantly higher for other (4.976 ± 0.963) than for the self (4.571 ± 1.035), *t* (41) = 3.460, *p_corr_* = 0.002, *p* = 0.001. Yet, no significant difference was observed between Self (1.367 ± 0.496) and Other (1.243 ± 0.477) in the Neutral condition, *t* (41) = −2.220, *p_corr_*= 0.064, *p* = 0.032. The results showed significant main effects for both Stimulus type, *F* (1,41) = 6.024, *p* = 0.018, η_p_^2^ = 0.128, and Facial expression, *F* (1,41) = 816.197, *p* < 0.001, η_p_^2^ = 0.952. Cognitive empathy was higher for Other (3.110 ± 2.024) than for Self (2.969 ± 1.802), and higher for Painful (4.774 ± 1.014) than for Neutral stimuli (1.305 ± 0.487).

### ERP results

#### N1

The mean N1 amplitude reached a maximum negative deflection in a time window of 130-190 ms (Figure 3). The results of the 2×2 within-subjects ANOVA (Stimulus type: Self vs. Other) × (Facial expression: Neutral vs. Painful) showed a non-significant interaction, *F* (1,41) = 0.422, *p* = 0.519, η_p_^2^=0.010. Similarly, the main effects of Stimulus type, *F* (1,41) = 2.143, *p* = 0.151, η_p_^2^ = 0.050, and Facial expression, *F* (1,41) = 1.103, *p* = 0.300, η_p_^2^ = 0.026, were not statistically significant.

**Figure 1.**
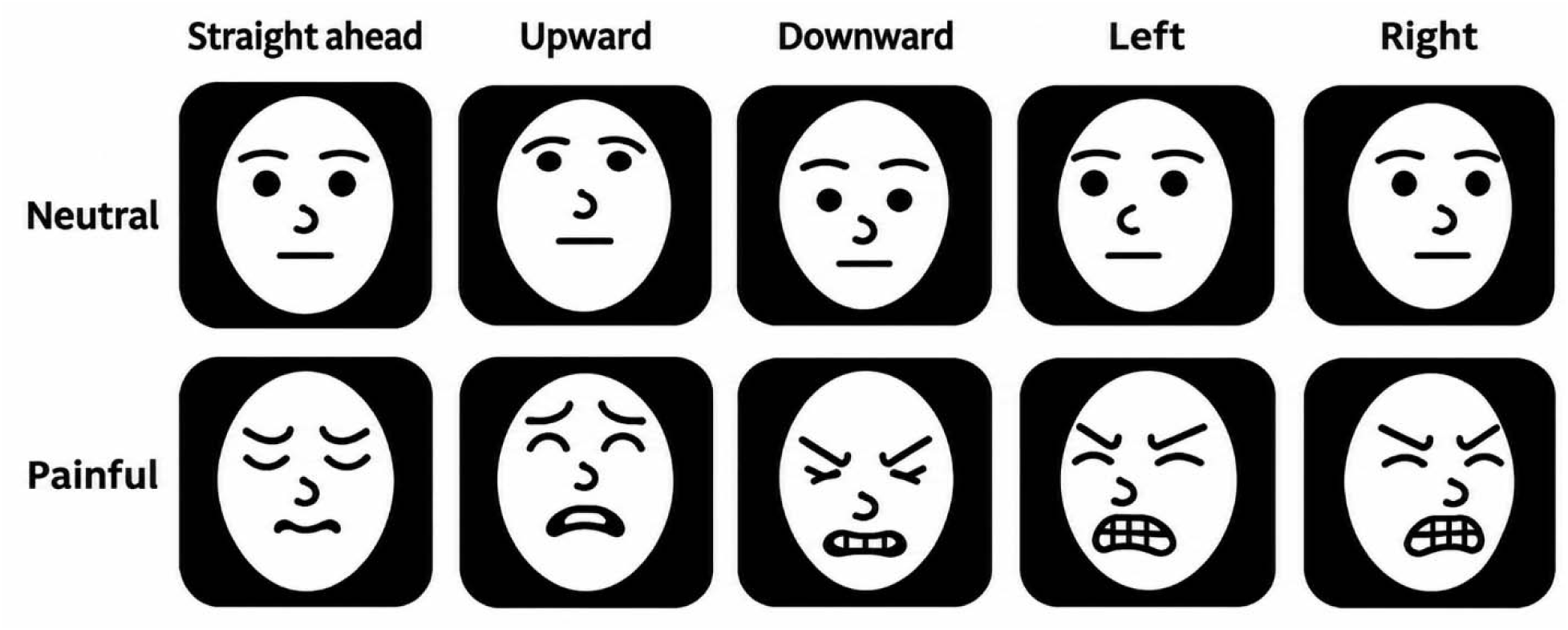
Example of stimuli used in the tasks.

**Figure 2.**
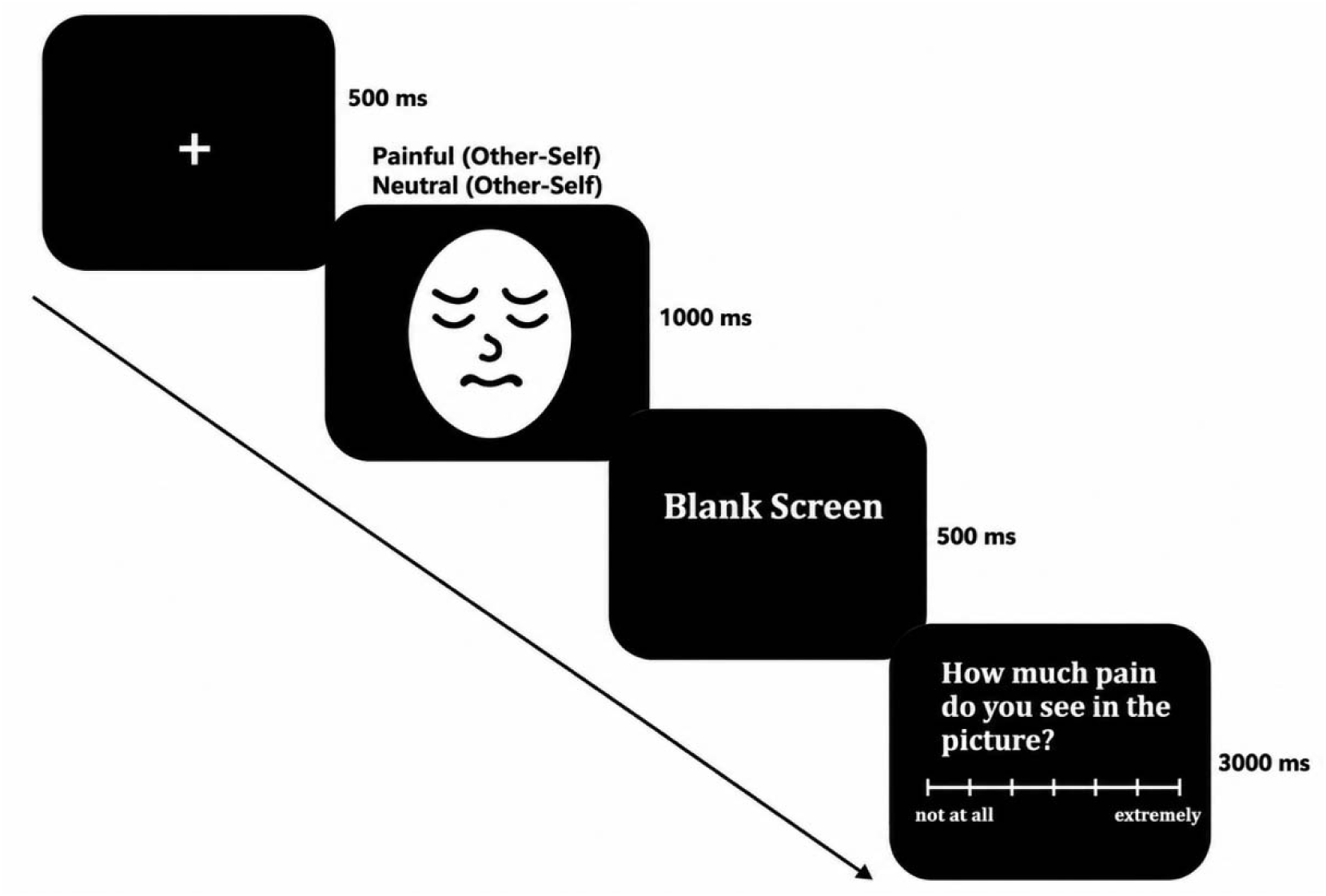
Flowchart Passive Viewing of Self and Other Painful Facial Expressions Task.

**Figure 3.**
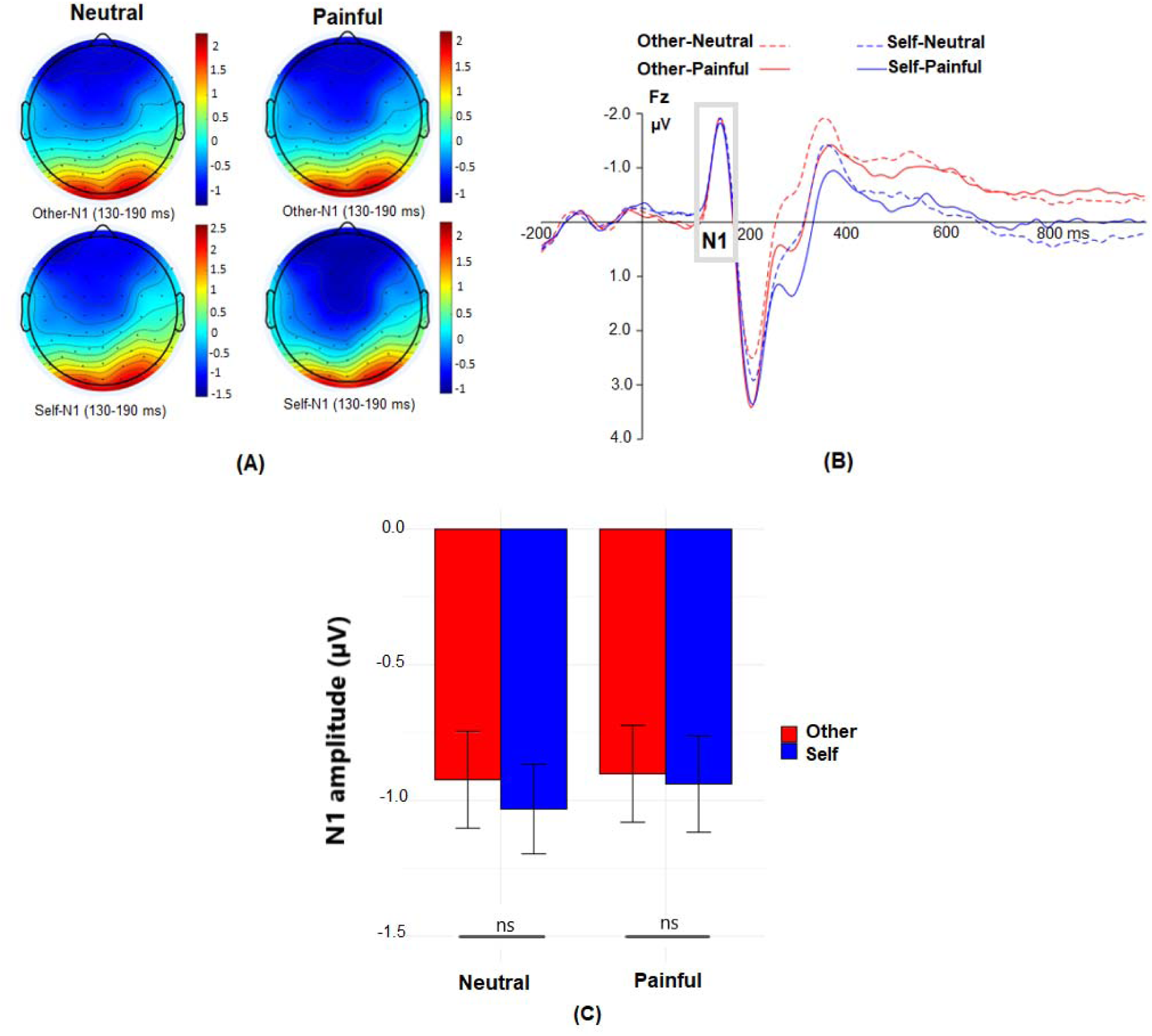
ERP results for the N1 component A) Topographic maps corresponding to the four experimental conditions at 130 to 190 ms. B) Grand-averaged ERPs of four experimental conditions recorded at the electrode Fz. C) Post hoc comparisons on Stimuli type × Facial expression interaction effects for N1 amplitude were conducted at the electrodes F1, Fz, F2, FC1, FCz, and FC2. Error bars represent standard errors of the mean.

#### N2

The mean N2 amplitude reached a maximum negative deflection in a time window of 170-290 ms. The ANOVA results showed a significant interaction between Stimulus type and Facial expression, *F* (1,41) = 5.729, *p* = 0.021, η_p_^2^ = 0.123. Post hoc comparisons revealed significant differences between Neutral and Painful stimuli in both the Other condition, *t* (41) = 6.314, *p_corr_* < 0.001, *p* < 0.001 (*M* _neutral_ = −1.202 ± 1.792; *M* _painful_ = −2.129 ± 2.400), and the Self condition, *t* (41) = 4.680, *p_corr_* <0.001, *p* < 0.001 (*M* _neutral_ = −1.906 ± 2.197; *M* _painful_ = −2.506 ± 2.643). However, the mean difference between Painful and Neutral stimuli was larger in the Other condition (mean difference = −0.927, SD = 0.951) than in the Self condition (mean difference = −0.600, SD = 0.831).

Additionally, Post hoc comparisons revealed significant differences between Other and Self stimuli in both the Neutral condition, *t* (41) = 5.698, *p_corr_* < 0.001, *p* < 0.001 (*M* _other_ = −1.202 ± 1.792; *M* _self_ = −1.906 ± 2.197), and the Painful condition, *t* (41) = 4.097, *p_corr_* < 0.001, *p* < 0.001 (*M* _other_ = −2.129 ± 2.400; *M* _self_ = −2.506 ± 2.643). However, the mean difference between Other and Self stimuli was larger in the Neutral condition (mean difference = −0.704, SD = 0.801) than in the Painful condition (mean difference = −0.377, SD = 0.597). The results showed significant main effects for both Stimulus type *F* (1,41) = 40.531, *p* < 0.001, η_p_^2^ = 0.497, and Facial expression, *F* (1,41) = 40.667; *p* < 0.001; η_p_^2^ = 0.498. The amplitude of N2 was more negative for Self (−2.206 ± 2.434) than for Other (−1.665± 2.156), and more negative for Painful (−2.317 ± 2.516) than for Neutral stimuli (−1.554 ± 2.024; Figure 4).

**Figure 4.**
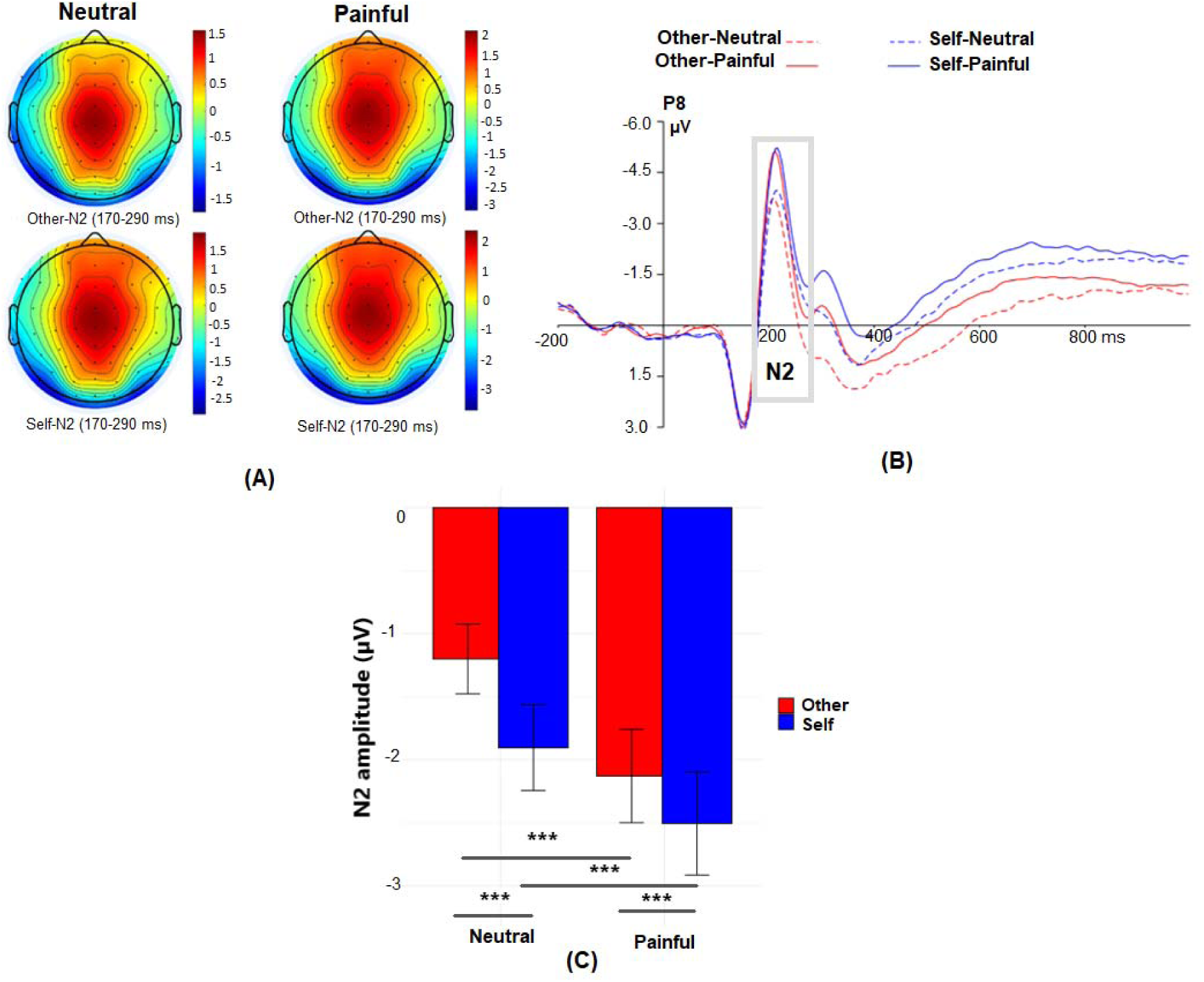
ERP results for the N2 component A) Topographic maps corresponding to the four experimental conditions at 170 to 290 ms. B) Grand-averaged ERPs of four experimental conditions recorded at the electrode P8. C) Post hoc comparisons on Stimuli type × Facial expression interaction effects for N2 amplitude were conducted at the electrodes P7, P8, PO7, and PO8. Error bars represent standard errors of the mean (\*\*\**p* < 0.001).

#### LPP

The mean LPP amplitude reached maximum positive deflection within the 340–800 ms time window. The ANOVA results showed a non-significant interaction between Stimulus type and Facial expression, *F* (1,41) = 0.169, *p* = 0.683, η_p_^2^ = 0.004. The ANOVA results showed statistically significant main effects for both Stimulus type, *F* (1,41) = 60.041, *p* < 0.001, η_p_^2^ = 0.594, and Facial expression, *F* (1,41) = 68.118, *p* < 0.001, η_p_^2^ = 0.624. The LPP amplitude was greater for Self (3.090 ± 1.620) than for Other (2.487 ± 1.434) and greater for Painful (3.264 ± 1.683) than for Neutral condition (2.312 ± 1.254; Figure 5).

**Figure 5.**
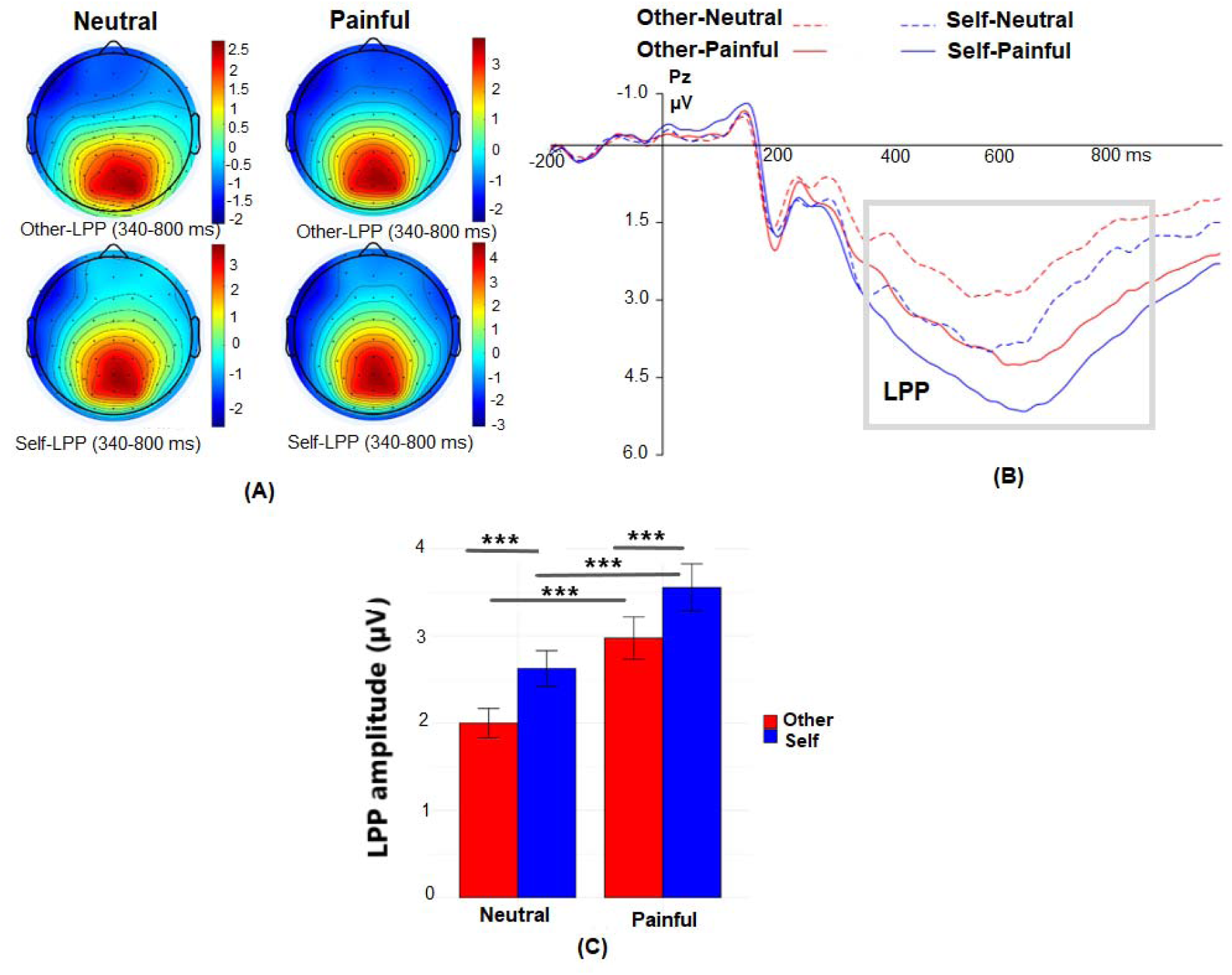
ERP results for the LPP component A) Topographic maps corresponding to the four experimental conditions at 340 to 800 ms. B) Grand-averaged ERPs of four experimental conditions recorded at the electrode Pz. C) Post hoc comparisons on Stimuli type × Facial expression interaction effects for LPP amplitude were conducted at the electrodes CP1, CPz, CP2, P1, Pz, P2, PO3, POz, and PO4 electrodes. Error bars represent standard errors of the mean (*** *p* < 0.001).

**Figure 6.**
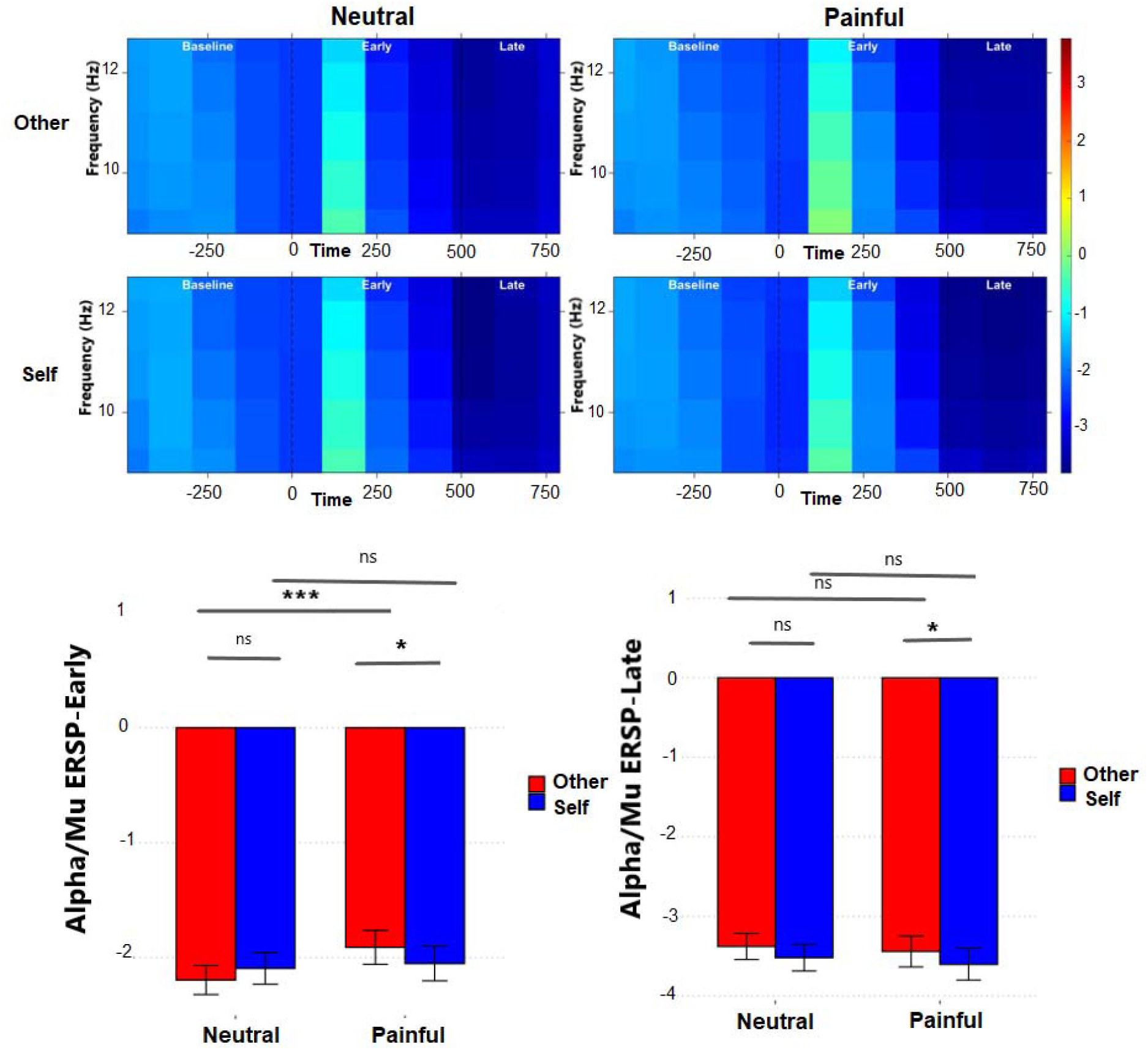
Time frequency results A) Alpha/Mu-ERD changes in sensorimotor regions (C3, Cz and C4) under different conditions. B) Post hoc comparisons of the Stimulus type × Facial expression interaction effect for Alpha/Mu ERD during the early and late time windows were conducted at electrodes C3, Cz, and C4. Error bars represent the standard error of the mean (\*\*\**p* < 0.001, * *p* < 0.05).

#### Time frequency analysis

The results of the three-way ANOVA revealed a significant interaction among Stimulus type (Self vs. Other), Facial expression (Painful vs. Neutral), and Time window (Early vs. Late) on Alpha/Mu rhythm oscillatory changes, *F* (1,41) = 4.676, *p* = 0.036, η_p_^2^ = 0.102. Main effects for both Stimulus type, *F* (1,41) = 4.661, *p* = 0.037, η_p_^2^ = 0.102, and Time window, *F* (1,41) = 146.805, *p* < 0.001, η_p_^2^ = 0.782, were also significant. However, the main effect of Facial expression was not statistically significant, *F* (1,41) = 0.797, *p* = 0.377, η_p_^2^ = 0.019. To further examine the pattern of the three-way interaction, the interaction between Stimulus type and Facial expression was examined separately within the Early and Late time windows. The results showed a significant interaction in the Early time window, *F* (1,41) = 9.024, *p* = 0.005, η_p_^2^ = 0.180, whereas no significant interaction was observed in the Late time window, *F* (1,41) = 0.043, *p* = 0.837, η_p_^2^ = 0.001. Given the significant interaction in the Early time window, simple effects were examined to further interpret this interaction. The results showed that, under Painful conditions, Alpha/Mu rhythm oscillatory changes differed significantly between Self (−2.051 ± 0.974) and Other stimuli (−1.910 ± 0.957), *t* (41) = 2.387, *p_corr_* = 0.022, *p* = 0.022, whereas this difference was not significant under Neutral conditions (Self: −2.093 ± 0.886; Other: −2.194 ± 0.814), *t* (41) = −1.577, *p_corr_*= 0.122, *p* = 0.122. Furthermore, for Other stimuli, Alpha/Mu rhythm oscillatory changes differed significantly between Painful (−1.190 ± 0.957) and Neutral conditions (−2.194 ± 0.814), *t* (41) = −4.391, *p_corr_*< 0.001, *p* < 0.001, whereas this difference was not significant for Self stimuli (Painful: −2.051± 0.974; Neutral: −2.093 ± 0.886), *t* (41) = −0.639, *p_corr_* = 0.526, *p* = 0.526.

## Discussion

The present study aimed to investigate the temporal dynamics of the neural mechanisms underlying empathic processing of pain in relation to the self and others. Participants viewed neutral and painful facial expressions of themselves and others while EEG was recorded. As expected, painful expressions elicited significantly higher cognitive and affective empathy compared to neutral expressions, indicating that the stimuli successfully engaged empathic processes. More importantly, our findings provide evidence that empathic processing is temporally dynamic and the contribution of self-other distinction varies across different stages of processing. In line with previous studies (Liu et al., 2023; Pan et al., 2023; Suzuki et al., 2015), painful expressions elicited a more negative N2 amplitude and a larger LPP than neutral expressions, suggesting the involvement of both early and later stages of neural processing. However, an interesting pattern emerged when the target of empathy was taken into account. At the N2 stage, self-related stimuli elicited more negative amplitudes than other-related stimuli in both neutral and painful conditions, indicating a consistent self–other distinction in early neural processing. Notably, the magnitude of this difference was greater for neutral than for painful expressions. One possible interpretation is that, in the absence of pain-related cues, self-related stimuli receive preferential processing due to their personal relevance. The presence of pain-related cues may heighten attention to pain-related information and engage empathic processes toward others’ pain, thereby potentially modulating the neural differentiation between self-related and other-related stimuli. Importantly, even in the context of painful expressions, self-related stimuli elicited more negative N2 amplitudes than other-related stimuli, suggesting that self-related processing remains particularly prominent during the early stages of empathic processing. In contrast, although the LPP amplitude were sensitive to both painfulness and self-relevance, the pain-related modulation of LPP did not differ between self and other. A similar temporal pattern emerged in the time-frequency domain, where during the early stages of processing, painful self-related expressions showed a higher suppression of Alpha/mu rhythm compared to another person. These together suggest that empathic response and shared representations are not temporally uniform processes but evolve across time, with self-other distinction influencing the earlier stages of empathic processing.

The more negative N2 amplitudes elicited by self-related painful expressions compared with other-related painful expressions indicate an early self–other distinction in pain processing. The N2 component has been implicated in early and automatic aspects of pain processing, affective sharing, and affective empathy (Liao et al., 2021; Pan et al., 2023; Suzuki et al., 2015; Vecchio & De Pascalis, 2023). Our findings therefore suggest that although shared representations may emerge relatively early during empathic processing, the strength of these responses is not equivalence for self- and other-related stimuli. Instead, early affective processing appears to be more strongly attuned to changes in the emotional state of the self. At the same time, the N2 has also been associated with face familiarity and the neural differentiation of self from other faces (Pierce et al., 2011). This raises the possibility that the observed N2 effects partly reflect enhanced processing of highly familiar self-related stimuli. However, the interaction between stimulus identity and painful expression suggests that familiarity or self-relevance alone cannot fully account for the observed pattern, as the magnitude of the self–other difference varied depending on whether the expressions were painful or neutral. Nevertheless, self-related stimuli elicited more negative N2 amplitudes than other-related stimuli in both conditions, indicating that early self–other differentiation was maintained even in the presence of pain-related cues. Rather than reflecting a mere familiarity effect, this pattern suggests that self-relevance may modulate the neural processing of pain-related information during relatively early stages of processing.

This pattern changed during later stages of processing. Painful stimuli elicited larger LPP amplitudes than neutral stimuli regardless of whether they were self-or other-related. At the same time, self-related stimuli elicited larger LPP amplitudes than other-related stimuli. Yet, the magnitude of the pain-related increase in LPP amplitude did not differ between self and other.

This finding is consistent with Brancadell et al. (2024), who reported larger LPP amplitudes for self-related than other-related stimuli regardless of whether those stimuli were painful or neutral. Such findings may reflect the sustained allocation of attentional and motivational resources to self-related information. Likewise, the larger LPP amplitudes observed for painful compared to neutral stimuli are consistent with previous findings showing enhanced late positive activity in response to pain-related information (Liu et al., 2023; Miedzobrodzka et al., 2023).

Together these findings indicated that later stages of processing remain sensitive both to the emotional significance of the stimulus and to its self-relevance, but that the modulation associated specifically with pain is no longer dependent on whether the stimulus represents the self or another person. One possibility is that, as empathic processing progresses, reliance on strongly self-oriented representations gradually decreases, allowing increasingly shared self– other representations to facilitate a more accurate understanding of another person’s emotional experience. Alternatively, later stages of empathic processing may depend more on cognitive appraisal and evaluation of the emotional significance of the stimulus, thereby reducing the relative contribution of self-other distinction. These interpretations are not mutually exclusive and suggest a transition from relatively self-dependent early processing toward more integrative and evaluative processing at later stages.

In contrast to N2 and LPP findings, as expected, we observed no significant modulation of N1 amplitude as a function of either pain condition or stimulus identity. This result is consistent with several previous studies reporting no reliable N1 differentiation between painful and neutral stimuli in empathy for pain paradigms (Ma et al., 2019; Sessa et al., 2014; Sheng & Han, 2012; Suzuki et al., 2015). Specifically, a meta-analysis of ERP studies similarly demonstrated that observing others in pain does not reliably modulate the N1 component (Coll, 2018), suggesting that the N1 may not constitute a sensitive or robust electrophysiological index of affective sharing. More generally, the absence of N1 modulation may indicate that the earliest stages of visual processing are primarily dominated by low-level perceptual and structural characteristics of the stimuli, including the initial structural encoding of faces (Balas & Conlin, 2015), before more elaborate differentiation based on self-relevance or emotional meaning emerges. This interpretation is also compatible with evidence suggesting limited differentiation between self and other faces during the very early stages of processing (Alzueta et al., 2019).

Interestingly, the temporal dynamics observed in the ERP components were paralleled by changes in the alpha/mu suppression. The interaction between stimulus identity and facial expression emerged in the early time window but was absent during the later time window, further suggesting that influence of self-other distinction is more pronounced during the early stages of empathy for pain and pain-related processing. Rather than reflecting a general enhancement of sensorimotor engagement with self-related pain, the present findings indicate that the processing of painful information depends on whether the observed experience is attributed to the self or to another person. Our findings are consistent with those of Riečanský et al. (2020), who showed that self-attribution can modulate mu rhythm suppression during pain observation. Accordingly, when a stimulus is evaluated as belonging to the self, greater engagement of sensorimotor systems may reflect the allocation of greater processing resources to self-related painful information and increased sensitivity to cues of bodily injury and potential threats to the body. The relatively reduced Alpha/Mu suppression for other-related painful stimuli, compared with neutral stimuli, during the early stages of processing may be interpreted as a regulatory mechanism that prevents excessive engagement of the observer with another person’s pain experience and associated personal distress (Mu et al., 2008).

Overall, the ERP and time–frequency findings converge in suggesting that self–other distinction plays a particularly important role during the early stages of empathy processing. Early empathic responses appear to be partly dominated by self-oriented representations, such that the processing of pain-related information is stronger when the observed experience is linked to the self. These results are compatible with theoretical accounts proposing that understanding another person’s pain is initially shaped by one’s own representations and personal experiences of pain (Decety & Hodges, 2006; Lamm et al., 2011). As processing progresses, however, the influence of self-other distinction on the pain-related neural responses appears to diminish, potentially reflecting a shift toward more abstract, integrative, and cognitively elaborated representations of emotional experience. Thus, rather than constituting a single unitary mechanism, empathy may emerge from a sequence of partially distinct processes in which self-related representations contribute differently across time.

Despite promising findings, our study also has limitations. The sample was homogeneous with respect to both gender and age. Participants were exclusively women between 18 and 34 years of age. This limits the generalizability of the findings to other age and gender groups.

Additionally, the use of simulated images instead of direct experience or induction of pain in painful situations may have influenced empathic responses. Future research should address the limitations of the present study by examining whether these findings generalize across male and female participants and across different age groups. In addition, future studies could investigate whether similar effects emerge when using stimuli depicting real pain experiences, rather than static images of individuals posing painful expressions while imagining pain. Beyond advancing the theoretical understanding of the neural basis of empathy, the present findings may also have important practical implications for understanding conditions in which empathic processing or self-other representation differs from that observed in the general population. Investigating the temporal organization of these mechanisms in populations characterized by impaired empathic processing, such as autistic individuals, may help clarify which stages of empathic processing are altered and whether such differences primarily involve early self-other mapping, later cognitive evaluation, or their interaction. More broadly, the present results highlight the importance of considering empathy as a temporally evolving process and demonstrate that the neural distinction between self and other is particularly pronounced during the relatively early stages of responding to pain-related information.

## Acknowledgments

The authors sincerely thank all participants for their valuable contributions to this study. We also thank the EEG laboratory at ICBS.

## Author contribution

**PR**: Conceptualization, Methodology, Data collection, Formal Analysis, Software, Writing- Original Draft, Writing- Review and Editing.**SG**: Methodology, Formal Analysis, Supervision, Writing-Original Draft, Writing- Review and Editing. **SHGH**: Conceptualization, Supervision, Writing- Review and Editing. **KB**: Conceptualization, Methodology, Supervision, Writing- Review and Editing.

## Data availability

The data collected during this study are available from the corresponding author upon reasonable request.

## Declaration of generative AI use

During the preparation of this manuscript, ChatGPT was used solely to assist with language editing and the correction of grammatical and stylistic issues. The authors reviewed and revised all suggested changes and take full responsibility for the content of the manuscript.

## References

Alzueta, E., Melcón, M., Poch, C., & Capilla, A. (2019). Is your own face more than a highly familiar face? Biol Psychol, 142, 100–107. 10.1016/j.biopsycho.2019.01.018

Balas, B., & Conlin, C. (2015). The visual N1 is sensitive to deviations from natural texture appearance. PLoS One, 10(9), e0136471.

Branchadell, V., Poy, R., Ribes-Guardiola, P., Segarra, P., & Moltó, J. (2024). Psychopathic callousness and perspective taking in pain processing: an ERP study. Soc Cogn Affect Neurosci, 19(1). 10.1093/scan/nsae022

Carretié, L., Hinojosa, J. A., Martín-Loeches, M., Mercado, F., & Tapia, M. (2004). Automatic attention to emotional stimuli: neural correlates. Hum Brain Mapp, 22(4), 290–299. 10.1002/hbm.20037

Chen, C., Yang, C. Y., & Cheng, Y. (2012). Sensorimotor resonance is an outcome but not a platform to anticipating harm to others. Soc Neurosci, 7(6), 578–590. 10.1080/17470919.2012.686924

Chen, T., Li, Q., Peng, M., & Li, X. (2023). Moral transgression modulates empathy for pain: Evidence from ERP and EEG data. Biol Psychol, 176, 108467. 10.1016/j.biopsycho.2022.108467

Chen, Y., Liang, J., Han, G., Yang, X., & Song, J. (2024). Peer Status Influences In-Group Favoritism in Pain Empathy During Middle Childhood: Evidence from Behavioral and Event-Related Potentials Studies. Brain Sci, 14(12). 10.3390/brainsci14121262

Cheng, Y., Chen, C., & Decety, J. (2014). An EEG/ERP investigation of the development of empathy in early and middle childhood. Dev Cogn Neurosci, 10, 160–169. 10.1016/j.dcn.2014.08.012

Coll, M. P. (2018). Meta-analysis of ERP investigations of pain empathy underlines methodological issues in ERP research. Soc Cogn Affect Neurosci, 13(10), 1003–1017. 10.1093/scan/nsy072

Coll, M. P., Bird, G., Catmur, C., & Press, C. (2015). Cross-modal repetition effects in the mu rhythm indicate tactile mirroring during action observation. Cortex, 63, 121–131. 10.1016/j.cortex.2014.08.024

Craig, K. D., Prkachin, K. M., & Grunau, R. V. (1992). The facial expression of pain. Handbook of pain assessment, 2, 257–276.

Cui, F., Zhu, X., Gu, R., & Luo, Y.-j. (2016). When your pain signifies my gain: neural activity while evaluating outcomes based on another person’s pain. Scientific Reports, 6(1), 26426. 10.1038/srep26426

Davis, M. H. (1983). Measuring individual differences in empathy: Evidence for a multidimensional approach. Journal of Personality and Social Psychology, 44(1), 113–126. 10.1037/0022-3514.44.1.113

Davis, M. H. (2006). Empathy. In J. E. Stets & J. H. Turner (Eds.), Handbook of the Sociology of Emotions (pp. 443–466). Springer US. 10.1007/978-0-387-30715-2_20

Decety, J., & Hodges, S. D. (2006). The social neuroscience of empathy. In Bridging social psychology (pp. 121–128). Psychology Press.

Decety, J., & Jackson, P. L. (2004). The functional architecture of human empathy. Behav Cogn Neurosci Rev, 3(2), 71–100. 10.1177/1534582304267187

Decety, J., & Lamm, C. (2006). Human Empathy Through the Lens of Social Neuroscience. TheScientificWorldJournal, 6, 1146–1163. 10.1100/tsw.2006.221

Decety, J., & Sommerville, J. A. (2003). Shared representations between self and other: a social cognitive neuroscience view. Trends Cogn Sci, 7(12), 527–533. 10.1016/j.tics.2003.10.004

Delorme, A., & Makeig, S. (2004). EEGLAB: an open source toolbox for analysis of single-trial EEG dynamics including independent component analysis. J Neurosci Methods, 134(1), 9–21. 10.1016/j.jneumeth.2003.10.009

Dvash, J., & Shamay-Tsoory, S. G. (2014). Theory of mind and empathy as multidimensional constructs: Neurological foundations. Topics in Language Disorders, 34(4), 282–295. 10.1097/TLD.0000000000000040

Ekman, P., & Friesen, W. V. (1978). Facial action coding system. Environmental psychology & nonverbal behavior.

Fabi, S., & Leuthold, H. (2017). Empathy for pain influences perceptual and motor processing: Evidence from response force, ERPs, and EEG oscillations. Social Neuroscience, 12(6), 701–716. 10.1080/17470919.2016.1238009

Fan, Y., & Han, S. (2008). Temporal dynamic of neural mechanisms involved in empathy for pain: an event-related brain potential study. Neuropsychologia, 46(1), 160–173. 10.1016/j.neuropsychologia.2007.07.023

Faul, F., Erdfelder, E., Lang, A. G., & Buchner, A. (2007). G*Power 3: a flexible statistical power analysis program for the social, behavioral, and biomedical sciences. Behav Res Methods, 39(2), 175–191. 10.3758/bf03193146

Ferguson, H. J., De Lillo, M., Woodrow-Hill, C., Foley, R., & Bradford, E. E. F. (2024). Neural empathy mechanisms are shared for physical and social pain, and increase from adolescence to older adulthood. Soc Cogn Affect Neurosci, 19(1). 10.1093/scan/nsae080

Gaetz, W., & Cheyne, D. (2006). Localization of sensorimotor cortical rhythms induced by tactile stimulation using spatially filtered MEG. Neuroimage, 30(3), 899–908. 10.1016/j.neuroimage.2005.10.009

Golbabaei, S., & Borhani, K. (2024). Nearsighted empathy: exploring the effect of empathy on distance perception, with eye movements as modulators. Scientific Reports, 14(1), 25146. 10.1038/s41598-024-76731-0

Golbabaei, S., Hosseinnezhad, M., Karami, Z., & Borhani, K. (2026). Empathy shortens perceived time in association with visual attention but not physiological measures. Scientific Reports, 16(1), 32300.

Golbabaei, S., Walter, M., & Borhani, K. (2026). Shared Pain, Shared Decisions: How Empathy Shapes Social Conformity Through Physiology and Visual Attention. bioRxiv, 2026.2006. 2027.734940.

Gomarus, H. K., Althaus, M., Wijers, A. A., & Minderaa, R. B. (2006). The effects of memory load and stimulus relevance on the EEG during a visual selective memory search task: an ERP and ERD/ERS study. Clin Neurophysiol, 117(4), 871–884. 10.1016/j.clinph.2005.12.008

Grice-Jackson, T., Critchley, H. D., Banissy, M. J., & Ward, J. (2017). Common and distinct neural mechanisms associated with the conscious experience of vicarious pain. Cortex, 94, 152–163. 10.1016/j.cortex.2017.06.015

Hajcak, G., & Foti, D. (2020). Significance?… Significance! Empirical, methodological, and theoretical connections between the late positive potential and P300 as neural responses to stimulus significance: An integrative review. Psychophysiology, 57(7), e13570.

Jackson, P. L., Brunet, E., Meltzoff, A. N., & Decety, J. (2006). Empathy examined through the neural mechanisms involved in imagining how I feel versus how you feel pain. Neuropsychologia, 44(5), 752–761. 10.1016/j.neuropsychologia.2005.07.015

Jackson, P. L., Meltzoff, A. N., & Decety, J. (2005). How do we perceive the pain of others? A window into the neural processes involved in empathy. Neuroimage, 24(3), 771–779. 10.1016/j.neuroimage.2004.09.006

Jung, T. P., Makeig, S., Westerfield, M., Townsend, J., Courchesne, E., & Sejnowski, T. J. (2001). Analysis and visualization of single-trial event-related potentials. Hum Brain Mapp, 14(3), 166–185. 10.1002/hbm.1050

Knoblich, G., & Flach, R. (2003). Action identity: evidence from self-recognition, prediction, and coordination. Conscious Cogn, 12(4), 620–632. 10.1016/s1053-8100(03)00070-9

Lamm, C., Bukowski, H., & Silani, G. (2016). From shared to distinct self-other representations in empathy: evidence from neurotypical function and socio-cognitive disorders. Philos Trans R Soc Lond B Biol Sci, 371(1686), 20150083. 10.1098/rstb.2015.0083

Lamm, C., Decety, J., & Singer, T. (2011). Meta-analytic evidence for common and distinct neural networks associated with directly experienced pain and empathy for pain. Neuroimage, 54(3), 2492–2502. 10.1016/j.neuroimage.2010.10.014

Lechinger, J., Wielek, T., Blume, C., Pichler, G., Michitsch, G., Donis, J., Gruber, W., & Schabus, M. (2016). Event-related EEG power modulations and phase connectivity indicate the focus of attention in an auditory own name paradigm. J Neurol, 263(8), 1530–1543. 10.1007/s00415-016-8150-z

Li, X., Liu, Y., Ye, Q., Lu, X., & Peng, W. (2020). The linkage between first-hand pain sensitivity and empathy for others’ pain: Attention matters. Hum Brain Mapp, 41(17), 4815–4828. 10.1002/hbm.25160

Li, X., Zhang, Y., Xiang, B., & Meng, J. (2019). Differences between empathy for face and body pain: Cognitive and neural responses. Brain Science Advances, 5(4), 256–264. 10.26599/BSA.2019.9050022

Li, Y., Wei, Z., Shao, M., Hong, M., Yang, D., Luo, L., & Meng, J. (2022). Empathy for pain in individuals with autistic traits during observation of static and dynamic stimuli. Front Psychiatry, 13, 1022087. 10.3389/fpsyt.2022.1022087

Liao, W., Zhang, Y., Huang, X., Xu, X., & Peng, X. (2021). “Emoji, I can feel your pain” - Neural responses to facial and emoji expressions of pain. Biol Psychol, 163, 108134. 10.1016/j.biopsycho.2021.108134

Liu, S., Tang, F., Dou, H., & Zhang, W. (2023). The relationship between autistic traits and empathy in adolescents: An ERP study. Neurosci Lett, 802, 137173. 10.1016/j.neulet.2023.137173

Liu, Y., Meng, J., Yao, M., Ye, Q., Fan, B., & Peng, W. (2019). Hearing other’s pain is associated with sensitivity to physical pain: An ERP study. Biol Psychol, 145, 150–158. 10.1016/j.biopsycho.2019.03.011

Ma, X., Wu, K., & Zhang, E. (2019). The Role of Social Power in Neural Responses to Others’ Pain. Front Psychol, 10, 2320. 10.3389/fpsyg.2019.02320

McDonald, B., & Kanske, P. (2023). Gender differences in empathy, compassion, and prosocial donations, but not theory of mind in a naturalistic social task. Sci Rep, 13(1), 20748. 10.1038/s41598-023-47747-9

Meng, J., Shen, L., Li, Z., & Peng, W. (2019). Top-down Effects on Empathy for Pain in Adults with Autistic Traits. Sci Rep, 9(1), 8022. 10.1038/s41598-019-44400-2

Mestre, M. V., Samper, P., Frías, M. D., & Tur, A. M. (2009). Are Women More Empathetic than Men? A Longitudinal Study in Adolescence. The Spanish journal of psychology, 12(1), 76–83. 10.1017/S1138741600001499

Miedzobrodzka, E., van Hooff, J. C., Krabbendam, L., & Konijn, E. A. (2023). Desensitized gamers? Violent video game exposure and empathy for pain in adolescents - an ERP study. Soc Neurosci, 18(6), 365–381. 10.1080/17470919.2023.2284999

Mu, Y., Fan, Y., Mao, L., & Han, S. (2008). Event-related theta and alpha oscillations mediate empathy for pain. Brain Res, 1234, 128–136. 10.1016/j.brainres.2008.07.113

Oberman, L. M., & Ramachandran, V. S. (2007). The simulating social mind: the role of the mirror neuron system and simulation in the social and communicative deficits of autism spectrum disorders. Psychol Bull, 133(2), 310–327. 10.1037/0033-2909.133.2.310

Olofsson, J. K., Nordin, S., Sequeira, H., & Polich, J. (2008). Affective picture processing: an integrative review of ERP findings. Biol Psychol, 77(3), 247–265. 10.1016/j.biopsycho.2007.11.006

Pan, Z., Zhang, C., Su, W., Qi, X., Feng, X., Gao, L., Xu, X., & Liu, J. (2023). Relationship between individual differences in pain empathy and task-and resting-state EEG. Neuroimage, 284, 120452. 10.1016/j.neuroimage.2023.120452

Peirce, J. W. (2007). PsychoPy--Psychophysics software in Python. J Neurosci Methods, 162(1-2), 8–13. 10.1016/j.jneumeth.2006.11.017

Perry, A., Bentin, S., Bartal, I. B., Lamm, C., & Decety, J. (2010). “Feeling” the pain of those who are different from us: Modulation of EEG in the mu/alpha range. Cogn Affect Behav Neurosci, 10(4), 493–504. 10.3758/cabn.10.4.493

Pfabigan, D. M., Seidel, E. M., Wucherer, A. M., Keckeis, K., Derntl, B., & Lamm, C. (2015). Affective empathy differs in male violent offenders with high-and low-trait psychopathy. J Pers Disord, 29(1), 42–61. 10.1521/pedi_2014_28_145

Pierce, L. J., Scott, L., Boddington, S., Droucker, D., Curran, T., & Tanaka, J. (2011). The N250 Brain Potential to Personally Familiar and Newly Learned Faces and Objects [Original Research]. *Frontiers in Human Neuroscience*, volume 5 *-*2011. 10.3389/fnhum.2011.00111

Riečanský, I., Lengersdorff, L. L., Pfabigan, D. M., & Lamm, C. (2020). Increasing self-other bodily overlap increases sensorimotor resonance to others’ pain. Cogn Affect Behav Neurosci, 20(1), 19–33. 10.3758/s13415-019-00724-0

Sessa, P., Meconi, F., Castelli, L., & Dell’Acqua, R. (2014). Taking one’s time in feeling other-race pain: an event-related potential investigation on the time-course of cross-racial empathy. Soc Cogn Affect Neurosci, 9(4), 454–463. 10.1093/scan/nst003

Sheng, F., & Han, S. (2012). Manipulations of cognitive strategies and intergroup relationships reduce the racial bias in empathic neural responses. Neuroimage, 61(4), 786–797. 10.1016/j.neuroimage.2012.04.028

Sun, Y. B., Lin, X. X., Ye, W., Wang, N., Wang, J. Y., & Luo, F. (2017). A Screening Mechanism Differentiating True from False Pain during Empathy. Sci Rep, 7(1), 11492. 10.1038/s41598-017-11963-x

Suzuki, Y., Galli, L., Ikeda, A., Itakura, S., & Kitazaki, M. (2015). Measuring empathy for human and robot hand pain using electroencephalography. Scientific Reports, 5(1), 15924. 10.1038/srep15924

Tajadura-Jiménez, A., Grehl, S., & Tsakiris, M. (2012). The other in me: interpersonal multisensory stimulation changes the mental representation of the self. PLoS One, 7(7), e40682. 10.1371/journal.pone.0040682

Vecchio, A., & De Pascalis, V. (2023). ERP indicators of situational empathy pain. Behavioural Brain Research, 439, 114224. 10.1016/j.bbr.2022.114224

Wang, H., Ao, L., Gao, Y., Liu, Y., & Zhang, X. (2023). Empathy for pain in individuals influenced by moral identity: Evidence from an ERP study. Physiol Behav, 266, 114202. 10.1016/j.physbeh.2023.114202

Woodruff, C. C., Martin, T., & Bilyk, N. (2011). Differences in self-and other-induced Mu suppression are correlated with empathic abilities. Brain Res, 1405, 69–76. 10.1016/j.brainres.2011.05.046

Yang, Y., Zhao, J., Zhang, H., Bi, T., Tian, J., Li, Q., & Guo, C. (2024). The mutual influences between working memory and empathy for pain: the role of social distance. Soc Cogn Affect Neurosci, 19(1). 10.1093/scan/nsae061

Zaki, J., Ochsner, K. N., Hanelin, J., Wager, T. D., & Mackey, S. C. (2007). Different circuits for different pain: patterns of functional connectivity reveal distinct networks for processing pain in self and others. Soc Neurosci, 2(3-4), 276–291. 10.1080/17470910701401973

Zhu, J., Wang, X. Q., He, X., Hu, Y. Y., Li, F., Liu, M. F., & Ye, B. (2019). Affective and Cognitive Empathy in Pre-teachers With Strong or Weak Professional Identity: An ERP Study. Front Hum Neurosci, 13, 175. 10.3389/fnhum.2019.00175

